# The UPR^mt^ Preserves Mitochondrial Import to Extend Lifespan

**DOI:** 10.1101/2020.07.01.182980

**Authors:** Nan Xin, Jenni Durieux, Chunxia Yang, Suzanne Wolff, Hyun-Eui Kim, Andrew Dillin

## Abstract

The mitochondrial unfolded protein response (UPR^mt^) is dedicated to promote mitochondrial proteostasis and is linked to extreme longevity in worms, flies, and mice. The key regulator of this process is the transcription factor, ATFS-1. In the absence of mitochondrial stress, ATFS-1 is transported to the mitochondria and degraded. During conditions of mitochondrial stress, ATFS-1 is excluded from the mitochondria and enters the nucleus to regulate the expression of UPR^mt^ genes. However, there exists a dichotomy in regards to induction of the UPR^mt^ and mitochondrial import. The repair proteins synthesized as a direct result of UPR^mt^ activation must be transported into damaged mitochondria that had previously excluded ATFS-1 due to reduced import efficiency. To address this conundrum, we analyzed the role of the import machinery under conditions where the UPR^mt^ was induced. Using *in vitro* biochemical assays of mitochondrial import and *in vivo* analysis of mitochondrial proteins, we surprisingly find that the efficiency of mitochondrial import increases when the UPR^mt^ is activated in an ATFS-1 dependent manner, even though membrane potential is reduced. The import machinery is upregulated at the transcription and translation level, and intact import machinery is essential for UPR^mt^-mediated increase and lifespan extension. With age, import capacity decreases, and activation of the UPR^mt^ delays this decline and increases longevity. Finally, we find that ATFS-1 has a significantly weaker mitochondrial targeting sequence (MTS), allowing for dynamic subcellular localization during the initial stages of UPR^mt^ activation.

## Introduction

During the course of eukaryotic evolution and the development of sequestered organelles, communication events co-evolved to allow proper homeostasis within the cell that encompassed master coordination by the nucleus. Such events have been discovered to include the unfolded protein response of the endoplasmic reticulum (UPR^ER^), the mitochondria (UPR^mt^), and the cytoplasmic heat shock response (HSR) ^1–3^. The primary mode of action of each of these stress pathways is the sensation of organelle-specific stress that is then communicated to the nucleus for transcriptional induction of the proper repair machinery to restore homeostasis within each organelle. Genes encoding compartment-specific chaperones are the sentinel transcriptional targets of such responses.

Mitochondria pose the most challenging organelle to coordinate stress responsiveness with the nucleus for several reasons. One, the mitochondrion is encapsulated within a double membrane system, the inner and outer membrane, thus creating a unique physical barrier that must be accommodated to signal from the mitochondrion to the nucleus. Two, the mitochondrion is composed of more than 1,000 different proteins, encoded within two distinct genomes, that comprise some of the largest complexes found in eukaryotic cells ^4,5^. Three, the by-products of oxidative respiration, superoxide radicals, pose a constant challenge to mitochondrial integrity ^6^. Four, import of proteins into the mitochondrion requires energy in the form of membrane potential and ATP created by the electron transport chain ^4^, and each cell can contain hundreds of mitochondria that fuse and divide at any given time to change the cellular mitochondrial landscape. Taken together, monitoring the integrity of the mitochondrion and all of its various protein complexes with the ability to communicate to the nucleus to ensure the integrity of this organelle is extremely complex.

Mitochondria can orchestrate and coordinate a wide number of different stress-response mechanisms under various cellular and subcellular perturbations. Such responses include the UPR^mt^, mitophagy, and programmed cell death. In response to mild mitochondrial stress, the UPR^mt^, a specific transcriptional stress response system that is mediated by ATFS-1, DVE-1, UBL-5, LIN-65, MET-2, and PHF-8, is activated to increase the production of mitochondrial localized chaperones and proteases to help relieve the stress ^7–11^. One of the major contributors to this response is ATFS-1. During mitochondrial stress, mitochondrial import efficiency is compromised, presumably due to depolarization of the mitochondrial membrane potential, which results in the inefficient import of the mitochondrial localized protein, ATFS-1. When ATFS-1 is not successfully imported into the mitochondria for degradation by mitochondrial proteases, it instead traffics to the nucleus, where it functions as a transcription factor, which coordinates with DVE-1, UBL-5 MET-2 and LIN-65 to induce the expression of mitochondrial chaperones and other genes required for repair of damaged mitochondria ^7–11^. Inherent within this model is the balance that must be maintained between the membrane potential, import machinery, and the ability to induce the UPR^mt^. However, the link between membrane potential, mitochondrial import, and the UPR^mt^ is largely unexplored.

With the current model of ATFS-1 localization dynamics during stress exists a dichotomy in regards to induction of the UPR^mt^ and mitochondrial import. The repair proteins synthesized as a direct result of UPR^mt^ activation by ATFS-1 must be transported into damaged mitochondria that must have a depolarized membrane potential preventing the import of ATFS-1 into the mitochondria, providing entry of ATFS-1 into the nucleus. If ATFS-1 is unable to enter the mitochondria during stress, how then are other proteins allowed entry, especially mitochondria with reduced membrane potential? Could there be coordination to increase mitochondrial import efficiency once the UPR^mt^ is induced? Is the mitochondrial import machinery a distinct branch of the UPR^mt^ to overcome the lack of integrity of damaged mitochondria? To address these questions, we analyzed the role of the import machinery under conditions where the UPR^mt^ was induced. Using *in vitro* biochemical assays of mitochondrial import and *in vivo* analysis of mitochondrially localized proteins, we find the efficiency of mitochondrial import increases when the UPR^mt^ is activated. More surprisingly, the increased import due to UPR^mt^ induction occurs when the mitochondrial membrane potential is decreased. Finally, we find that the induction of the import machinery is essential for UPR^mt^-mediated lifespan extension, and ectopic induction of the UPR^mt^ preserves import into late life.

## Results

### Assessing mitochondrial import capacity from *C. elegans* mitochondria

All but a few mitochondrial proteins are transcribed from the nuclear genome and imported post-translationally through the translocase of the outer/inner membrane (TOM/TIM) complex ^4,12^. Mitochondrial protein import depends on the mitochondrial membrane potential (ΔΨ), ATP, and is under the direction of mitochondrial targeting sequences (MTS), which is cleaved after import ^13^. To measure the efficiency of mitochondrial import, we adapted and validated a method in which substrate proteins are synthesized in an *in vitro* transcription/translation reaction, and subsequently imported into isolated mitochondria ^14,15^ (Fig. 1a). We used a model import substrate, su9-DHFR, in which the MTS of subunit 9 of the mitochondrial ATP synthase from *Neurospora* is fused to a fragment of the cytosolic protein dihydrofolate reductase (DHFR) from mice ^16^. An ATP regeneration system^17^ was applied to increase the efficiency of import. Importantly, a DHFR antibody can readily detect the fusion protein being imported (Fig. 1b, c, Extended Data Fig. 1a), as indicated by: 1) the change in size from a precursor protein to a mature DHFR; 2) the absence of mature DHFR upon disruption of membrane potential (ΔΨ); 3) the absence of precursor protein upon proteinase K treatment; and 4) the accumulation of a mature DHFR in a time-dependent manner. Mitochondrial import efficiency of su9-DHFR was reduced by 30-40% from mitochondrial preparations isolated from animals upon knocking down essential components of the TOM/TIM complex, *tomm-20* or *timm-17*, via RNAi, further validating the sensitivity and fidelity of the assay (Fig. 1d, e, Extended Data Fig. 1b).

**Figure 1.**
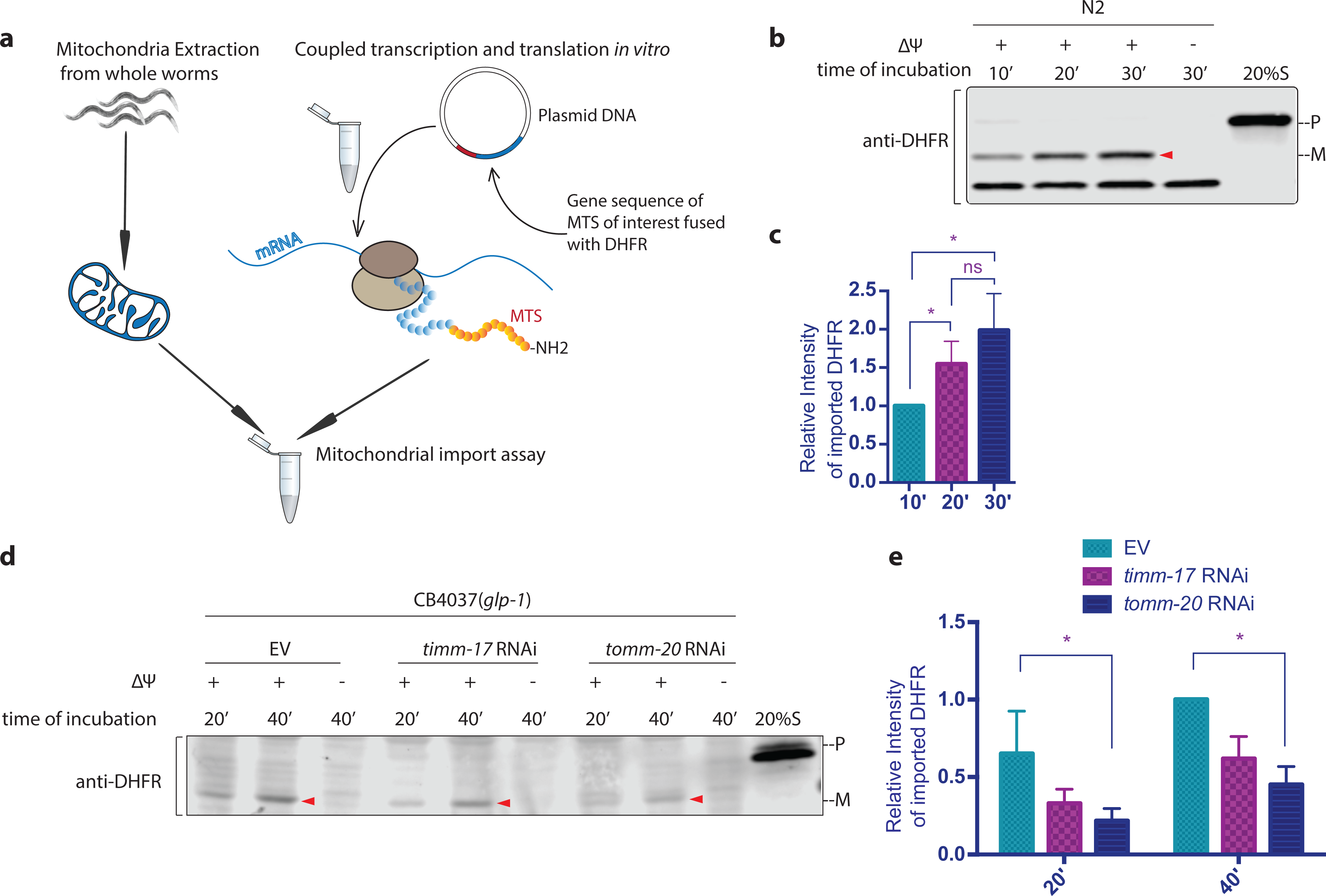
*C. elegans* mitochondrial protein *in vitro* import assay. **a**, Schematic diagram of the *C. elegans in vitro* mitochondrial protein import assay. **b** and **c**, su9-DHFR was transcribed and translated in a single reaction with the Quick Coupled Transcription/Translation System (TnT reaction). Mitochondria extraction was made from synchronized N2 wild type worms at day 1 of adulthood and quantified with BCA analysis. 50ug mitochondrial protein was used in each reaction. The substrate protein was incubated with mitochondria extraction in import buffer containing an ATP regeneration system for 10, 20, or 30 minutes at 25°C. Mitochondria were subsequently treated with proteinase K to remove non-imported proteins. Upon being imported, the MTS of su9 is cleaved. 2ug/ml valinomycin was used to disrupt the membrane potential (ΔΨ), thus inhibiting import. The precursor (p) and mature protein(m) were detected with the DHFR antibody by western blot analysis. Right lane: 20% of the su9-DHFR substrate used in the import assay representing the precursor (p). **d** and **e**, Germline-deficient, mutant *glp-1(e2141ts)* worms were bleach synchronized, grown at the restrictive temperature, 25°C, and treated with RNAi against *tomm-20* or *timm-17* until the first day of adulthood. Control worms were grown on bacteria containing empty vector alone. Mitochondria were isolated and subjected to the import assay. **c** and **e**, The efficiency of mitochondrial import was quantified by measuring the mature imported protein as detected by the DHFR antibody and analyzed with unpaired student’s t-test. All graphs are presented as mean ± SD of two or more biological repeats. *P<0.05. Arrowheads: mature (imported) DHFR with the MTS cleaved off.

### A critical difference in import capacity among germ and somatic cells in *C. elegans*

To determine whether induction of the UPR^mt^ had any impact upon mitochondrial import efficiency, we first induced the UPR^mt^ in wild-type N2 worms with RNAi against cytochrome c oxidase-1 subunit (*cco-1/cox-5B*), a component of the electron transport chain complex IV, and found that import efficiency was reduced in mitochondrial preparations isolated from whole animals (Extended Data Fig. 1c-e). This finding appears to be consistent with the current model of UPR^mt^ induction and ATFS-1 exclusion. However, we must note that worms treated with *cco-1* RNAi have reduced fecundity, notably due to a significantly underdeveloped germline ^18^. The soma of *C. elegans* is post-mitotic, containing only 959 cells; however, the germline is expansive and mitotic, containing both syncytial and cellularized cells. In fact, the development of the female germline accounts for the vast majority of mtDNA amplification in *C. elegans* ^19^. As the germline develops, rapid mitochondrial expansion is essential, and this could be a major component of the mitochondrial import activity found when entire animals are used to isolate the mitochondria; hence the reduced import efficiency of UPR^mt^ in animals treated with *cco-1* RNAi could be explained by the reduced germline found in these animals.

We, therefore, tested whether germline cells differ in import competency relative to post-mitotic cells. To this end, we used three temperature-sensitive sterile strains that differentially affect the development of the germline at the restrictive temperature. In particular, CB4037 *glp-1(e2141ts)* ^20^ and SS104 *glp-4(bn2ts)* ^21^ are sterile strains that lack the majority of the germline, whereas CF512 *fer-15(b26ts); fem-1(hc17)* ^22^ is sterile due to the conversion of sperm into oocytes. Of note, the sterile strain CF512 was used as the control to ensure that any difference we observed was not due to the contribution of import activity from the offspring. Comparison of the import of substrates into mitochondria isolated from the three sterile strains, we found that the germline-deficient mutants (*glp-1* and *glp-4*) were strikingly less import-competent than the sperm-deficient, oocyte proficient, CF512 strain, at the restrictive temperature (Extended Data Fig. 1f-h). In particular, the *glp-1* strain CB4037 is 80% lower than strain CF512 in import. In contrast, we observed no significant difference between these strains when raised at the permissive temperature, 15°C (Extended Data Fig. 1i-k), which allows normal germline development. Additionally, temporal shifting of *glp-1(e1241ts)* mutant animals from 15°C to 25°C during development allows limited germline development, with earlier shifts resulting in fewer germ cells, and later shifts having a near-complete complement of germ cells. By shifting at a series of time points, we found that import capacity was positively correlated with the number of germline cells present in the animals (Extended Data Fig. 1l-n). In contrast, the CF512 strain, which has a normal female germline at both permissive and restrictive temperatures, shows higher import competency at 25°C (Extended Data Fig. 1o-q). These results suggest that germline mitochondria are highly import-competent in comparison to post-mitotic, somatic cells in *C. elegans*.

### Mitochondrial import is enhanced upon UPR^mt^ induction

To understand if and how mitochondrial import is regulated upon mitochondrial stress in somatic cells, we induced UPR^mt^ in *C. elegans* and examined the import capacity of isolated mitochondria from animals lacking a germline. Considering the remarkable discrepancy in import capacity we found between somatic tissue and the germline, we genetically ablated the germline using *glp-1(e2141ts)* mutant animals ^18^, to only assay mitochondria from somatic tissues where the UPR^mt^ impacts longevity and health ^23,24^. We found that mitochondria isolated from *glp-1(e2141ts)* mutant animals treated with *cco-1* RNAi, import capacity was elevated 2 to 2.5 times that of age-matched, mock RNAi control treated animals (Fig. 2a, b, Extended Data Fig. 2a).

**Figure 2.**
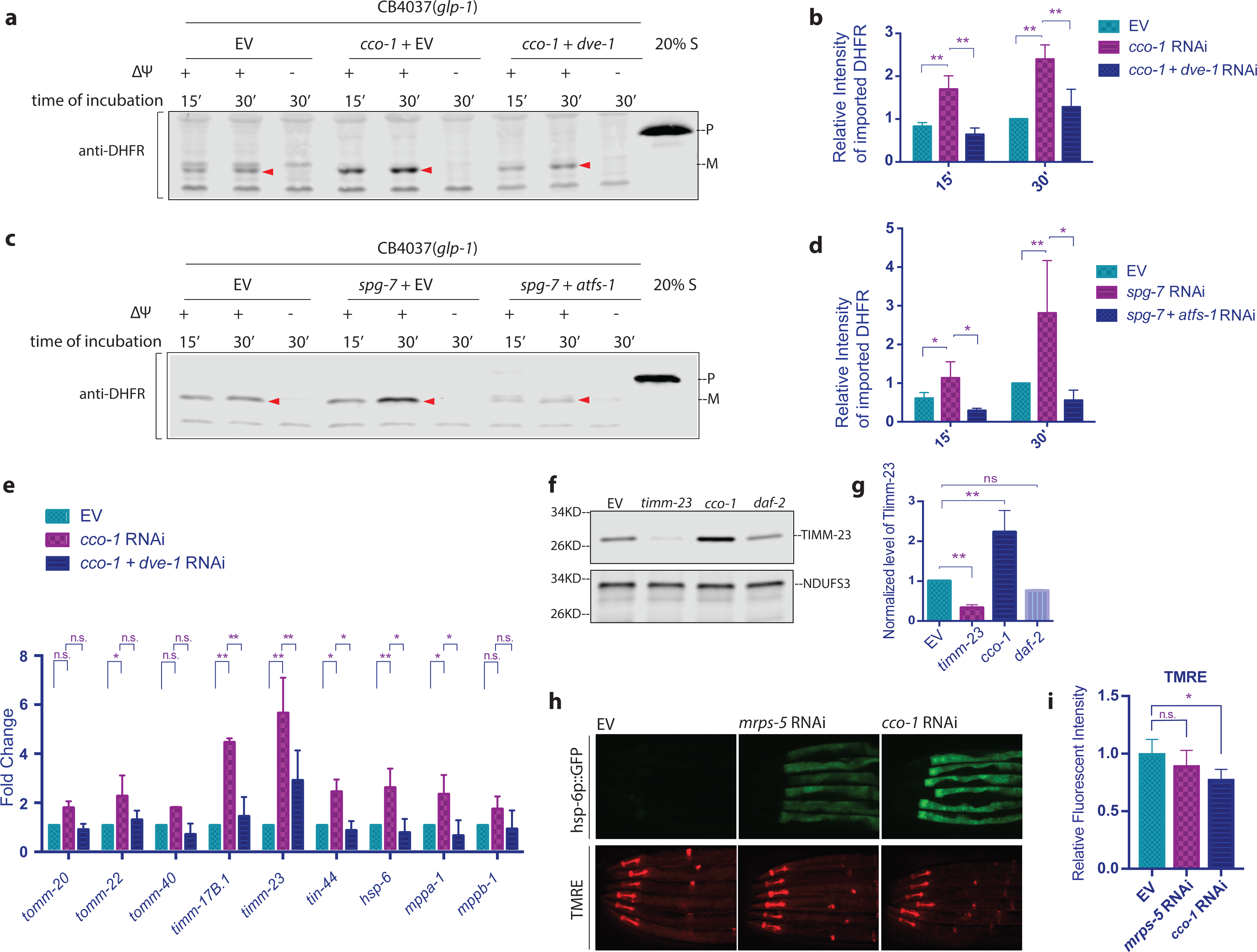
The UPR^mt^ promotes mitochondrial import. **a-e**, To induce UPR^mt^ during development, *glp-1(e2141ts*) mutant animals were grown at 25°C on bacteria expressing *cco-1* dsRNA (**a**,**b**,**e**) or *spg-7* dsRNA (**c**,**d**) from the time of hatching until the first day of adulthood (animals were treated with 1:1 mixture of bacteria containing the empty RNAi vector alone (EV) to match with the double RNAi treatment). To suppress UPR^mt^, animals were treated with double RNAi (1:1 mixture of bacteria replacing EV with bacteria expressing *dve-1* dsRNA (**a**,**b**,**e**) or *atfs-1* dsRNA (**c**,**d**)). Control worms were grown on bacteria containing empty vector (EV) alone. Mitochondria were isolated from the animals at day 1 of adulthood and subjected to the import assay followed by western blot analysis (**a, c**). Import efficiency was quantified by measuring the mature imported protein as detected by the DHFR antibody, followed by analysis with unpaired student’s t-test (**b, d**). **e**, RNA was isolated on day 1 of adulthood, and qPCR analysis was performed. **f** and **g**, *glp-1* animals were grown at 25°C on bacteria expressing *timm-23, cco-1*, or *daf-2* dsRNA (each was diluted to 1:1 ratio with bacteria containing the empty RNAi vector alone) from hatching until the first day of adulthood. Control worms were grown on bacteria containing the empty vector alone. Quantification is shown in **g** with unpaired student’s t-test. The signal intensity of TIMM-23 was normalized to that of NDUFS3. **h** and **i**, *hsp-6p*::GFP animals were grown at 20°C and treated with RNAi or empty RNAi vector control from hatching until the L4 stage, then transferred to plates of the same RNAi treatment with the addition of TMRE and grown overnight. Bacteria expressing *cco-1* dsRNA (right) or *mrps-5* dsRNA (middle) (both were diluted 20% with bacteria containing the empty RNAi vector alone) were used to induced UPR^mt^. Control worms were grown on bacteria containing the RNAi vector alone (left). Fluorescent intensity was quantified with Image J. All graphs are presented as mean ± SD of three or more biological repeats. *P<0.05, **P<0.01. Arrowheads: mature (imported) DHFR with the MTS cleaved off.

Struck by the robust increase in mitochondrial import efficiency conferred by *cco-1* RNAi in animals composed only of somatic cells, we next asked whether the increased import efficiency was a common response to mitochondrial stress. SPG-7 is an AAA protease involved in quality control of mitochondrial membrane proteins, as well as the assembly of protein complexes on the mitochondrial inner membrane ^25^. MRPS-5 is a mitochondrial ribosome protein ^23^. Knockdown of either *spg-7* or *mrps-5*, via RNAi, induces the UPR^mt^ and leads to lifespan extension of worms composed of post-mitotic, somatic cells ^23,26^. We found that mitochondrial import was also significantly enhanced upon either *spg-7* RNAi (Fig. 2c, d, Extended Data Fig. 2b) or *mrps-5* RNAi (Extended Data Fig. 2c-e). Taken together, both results suggest that the activation of the UPR^mt^ is widely associated with enhanced mitochondrial import.

To test whether the enhancement of import depends on the activation of UPR^mt^, we introduced *dve-1* RNAi into animals together with *cco-1* RNAi. DVE-1 is a transcription factor that mediates the activation of stress responsive genes upon UPR^mt 10^. *dve-1* RNAi partially suppressed the induction of *hsp-6p::gfp* reporter ^26^, indicating that knockdown of *dve-1* partially blocks the downstream effect of UPR^mt^ induced by *cco-1* RNAi (Extended Data Fig. 2f). We found that knockdown of *dve-1* resulted in marked suppression of import capacity that had been enhanced by *cco-1* knockdown (Fig. 2a, b), suggesting that the induction of the UPR^mt^ is essential for increased import efficiency. Importantly, consistent with earlier findings ^7,27^, ATFS-1 is required for the induction of UPR^mt^, as indicated by the *hsp-6p::gfp* reporter (Extended Data Fig. 2g). When *atfs-1* RNAi was introduced into animals together with *spg-7* RNAi, the enhancement of import was also suppressed (Fig. 2c, d). Taken together, these results indicate that the UPR^mt^ indeed mediates the upregulation of mitochondrial import.

### Mitochondrial import machinery is upregulated upon UPR^mt^ induction

The UPR^mt^ promotes mitochondrial protein homeostasis through signaling to the nucleus to induce the transcription of mitochondrial localized stress-responsive genes. Most of the protein products of the upregulated genes must then be imported into mitochondria to restore proteostasis. The proteins that constitute the import machinery, TIM/TOM complexes, are exclusively encoded by nuclear genes. Therefore, it is conceivable that the UPR^mt^ may enhance import through upregulating the expression of the TIM/TOM complex components. Indeed, the import machinery components *timm-17* and *timm-23* were upregulated upon UPR^mt^ induction by *spg-7* RNAi ^7^. Similarly, *timm-23* was also found to be moderately upregulated upon *cco-1* RNAi treatment ^8^. However, when we examined the RNA-sequencing data of animals treated with *cco-1* RNAi, we found no significant change in the transcription of most other import machinery components ^8,9^. As N2 worms were used in this RNA-seq analysis, the loss of germline caused by *cco-1* knockdown during development may counteract any effect in somatic mitochondria caused by *cco-1* RNAi. To examine the transcriptional regulation of import machinery in somatic tissue, we tested germline-deficient *glp-1(e2141ts)* mutant animals for the transcription of a series of genes encoding the mitochondrial import machinery. Comparing the synchronized and age-matched worms, we found that transcription of the TIM/TOM genes upon *cco-1* RNAi treatment was consistently higher than the mock RNAi control. Genes encoding TOM complex proteins, including *tomm-20, tomm-22*, and *tomm-40*, were enhanced one to two-fold. Core components of the TIM complex, *timm-17* and *timm-23*, were upregulated 3 and 7-fold, respectively (Fig. 2e). The transcription level of *timm-17* and *timm-23* at steady state appeared to be lower than other TIM/TOM component, whereas, their transcription was elevated to levels higher than other components upon UPR^mt^ activation. This is consistent with previous findings, which suggests that TIM23 protein might be the rate-limiting factor in mitochondrial import ^28^. Following the passage through the inner membrane pore formed by TIM17 and TIM23, the precursor proteins are pulled into the matrix by TIM44 and mtHSP70 (HSP-6). mtHSP70 also facilitates the proper folding of imported proteins, which are subsequently processed by MPP proteins, proteases that cleave off the mitochondrial targeting sequences (MTS) ^29^. We found that *hsp-6, tin-44*, as well as *mppa-1* and *mppb-1*, which are *C. elegans* homologs of mammalian *mtHsp70, Tim44*, and *Mpp*, respectively, were also upregulated by *cco-1* RNAi. Therefore, the entire repertoire of the import machinery appears to be transcriptionally induced by activation of the UPR^mt^ in somatic cells. Similarly, the TIM/TOM import machinery was also upregulated upon *spg-7* RNAi (Extended Data Fig. 2h). Additionally, the *cco-1* induced upregulation of import machinery genes was suppressed by RNAi against *dve-1* (Fig. 2e).

To further verify the regulation of import machinery, we generated an antibody against the *C. elegans* mitochondrial translocase protein TIMM-23 and verified its specificity with RNAi knockdown of *timm-23*. When comparing the endogenous level of TIMM-23, we found that the TIMM-23 protein level is indeed enhanced when the UPR^mt^ is activated (Fig. 2f, g).

Given that mitochondrial protein import requires membrane potential (ΔΨ), we examined if ΔΨ is enhanced upon UPR^mt^ activation. Surprisingly, in early Day 1 adult, the same stage when enhanced mitochondrial import was detected, we did not observe stronger TMRE staining, a marker of ΔΨ, in worms with activated UPR^mt^ (Fig. 2h, i). On the contrary, membrane potential was found to be reversely correlated with the activation of the UPR^mt^. In particular, knockdown of *cco-1* induced the UPR^mt^ more robustly than knockdown of *mrps-5*, as indicated by the level of *hsp-6p::gfp* reporter, whereas membrane potential is lower with *cco-1* knockdown (Fig. 2h, i). This is consistent with the model that mitochondrial stress leads to membrane depolarization. Together, these findings suggest that the enhancement of mitochondrial import we observed in the *in vitro* import assay is regulated through increased transcriptional regulation of import machinery, rather than enhanced membrane potential. Furthermore, and most surprisingly, these results indicate that aspects of mitochondrial import can be decoupled from membrane potential.

### The mitochondrial targeting sequence of ATFS-1 is less import competent

Presumably, the enhancement of import would allow efficient translocation of the repair proteins into the mitochondria to restore their proper function. However, previous findings suggest that import is compromised upon mitochondrial stress, thereby allowing ATFS-1 to translocate into the nucleus, where it activates the transcription of stress response genes ^2^. Is the import of ATFS-1 differentially regulated? To interrogate this possibility, we made a chimeric protein with the predicted mitochondrial targeting sequence of ATFS-1 (N terminal 73 amino acids) ^7^ fused with DHFR (ATFS1-DHFR). We found that upon the induction of the UPR^mt^, the import of ATFS1-DHFR is also upregulated (Fig. 3a, b, Extended Data Fig. 3a). Though the regulation of ATFS-1 import shows the same trend, we observed that the import of ATFS1-DHFR is less robust as compared to su9-DHFR (Fig. 3a). Indeed, when we compared the import of the two fusion proteins side by side, we found that mitochondrial import directed by the MTS of ATFS-1 is significantly less robust (Fig. 3a, c, Extended Data Fig. 3a). Our finding is consistent with the model that ATFS-1 has a weak MTS, which allows it to sense modest mitochondrial dysfunction and membrane depolarization to send a stress signal to the nucleus by translocation ^30^. Point mutations in the MTS of *atfs-1*, such as *et15* or *et18*, display constitutively active UPR^mt 33^, presumably due to lack of mitochondrial import of ATFS-1 and relocation to the nucleus. We introduced the *et15* and *et18* mutations into the MTS of the ATFS1-DHFR construct and used them in the *in vitro* import assay. We found that the mitochondrial targeting capacity deteriorated with both mutations in the MTS of *atfs-1* (Fig. 3d, Extended Data Fig. 3b).

**Figure 3.**
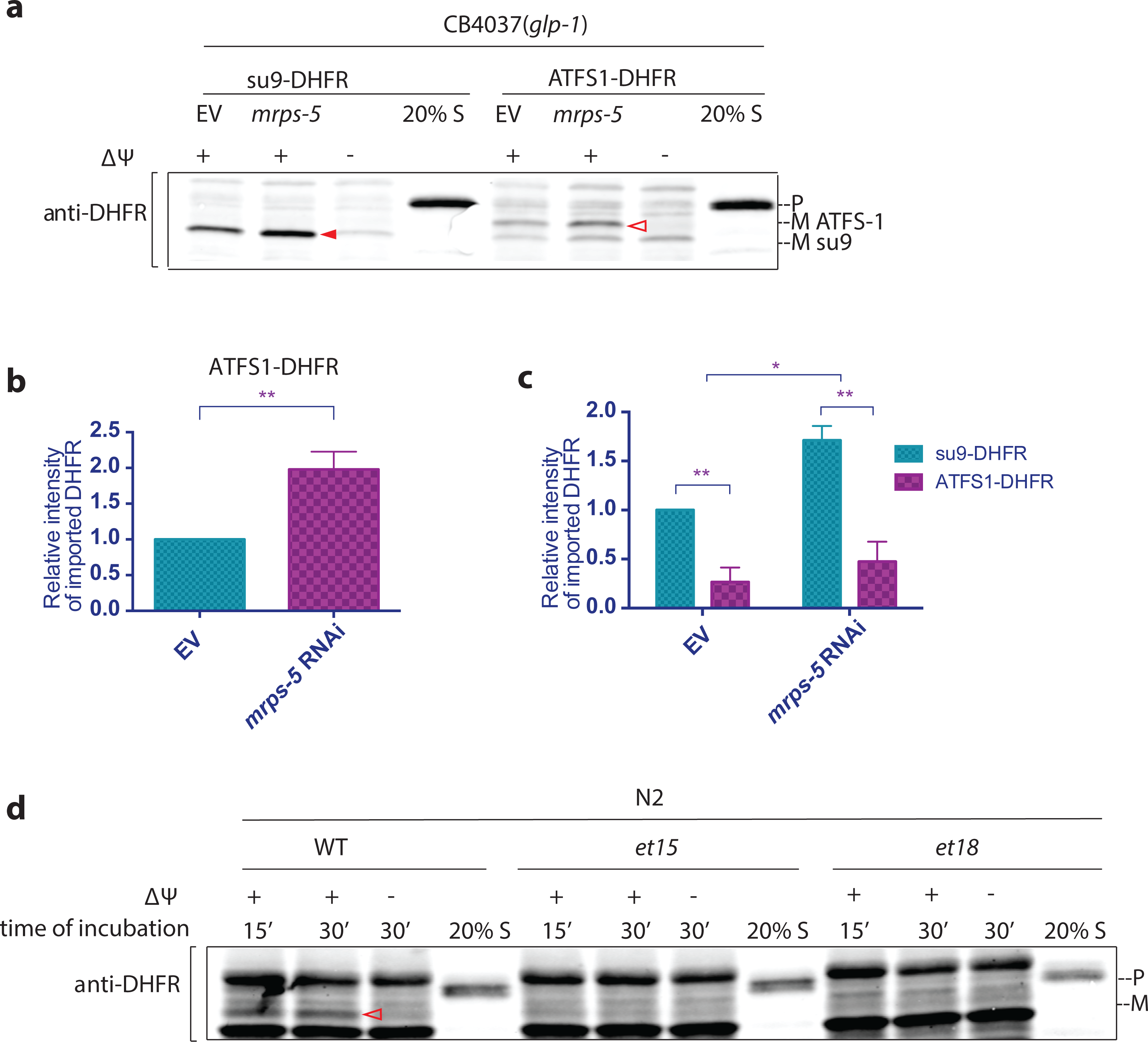
The Mitochondrial targeting sequence of ATFS-1 is less import competent. **a-c**, Comparison of import competency between su9-DHFR and ATFS-1-DHFR. The N-terminus 73 amino acid of ATFS-1 was used as MTS. *glp-1(e2141ts*) animals were grown at 25°C on bacteria expressing *mrps-5* dsRNA (20% diluted with bacteria containing the empty RNAi vector alone) from hatching until the first day of adulthood. Mitochondria were isolated on day 1 of adulthood and subjected to import assay with 30 minutes incubation time, followed by western blot analysis. Import efficiency was quantified by measuring the mature imported protein as detected by the DHFR antibody. Quantification of imported DHFR is shown in **b** and **c. b**, Import of ATFS1-DHFR was compared between *mrps-5* RNAi and empty vector control with unpaired student’s t-test. The graph is presented as mean ± SD of four biological repeats (*P<0.05, **P<0.01). **c**, Import of DHFR with the MTS of su9 and ATFS-1 were compared with two-way ANOVA. No interaction was found. The difference in MTS and the treatment with *mrps-5* RNAi both have significant effects on the import of DHFR. The graph is presented as mean ± SD of two biological repeats. *P<0.05, **P<0.01. **d**, Mitochondrial targeting capacity abolished by point mutations *et15* or *et18* in the MTS of ATFS-1. Mitochondria extraction was made from synchronized N2 wild type worms at day 1 of adulthood and subjected to import assay with different substrates. Import with *et15* or *et18* was below the detectable level. Solid arrowheads: mature (imported) su9-DHFR with the su-9 MTS cleaved off. Open arrowheads: mature (imported) ATFS1-DHFR with the MTS of ATFS-1 cleaved off.

### The lifespan extension caused by UPR^mt^ induction requires intact TIM/TOM complexes

Our findings suggest that the mitochondrial import machinery is upregulated at the transcriptional level upon UPR^mt^ induction, thereby elevating import capacity and allowing stress-responsive proteins to translocate into the mitochondrial matrix and restore proteostasis. One such stress-responsive proteins is the mitochondrial chaperone mtHSP70(HSP-6), which is transcriptionally upregulated upon the activation of UPR^mt^. Analyzing the subcellular fractionation of mitochondria from UPR^mt^ induced animals, we find increased levels of HSP-6 in the mitochondrial fraction (Fig. 4a, b, extended data Fig. 4a), suggesting that import *in vivo* is maintained at a level that allows efficient mitochondrial translocation of the elevated level of stress-responsive proteins, despite the reduction of membrane potential (Fig. 2h, i).

**Figure 4.**
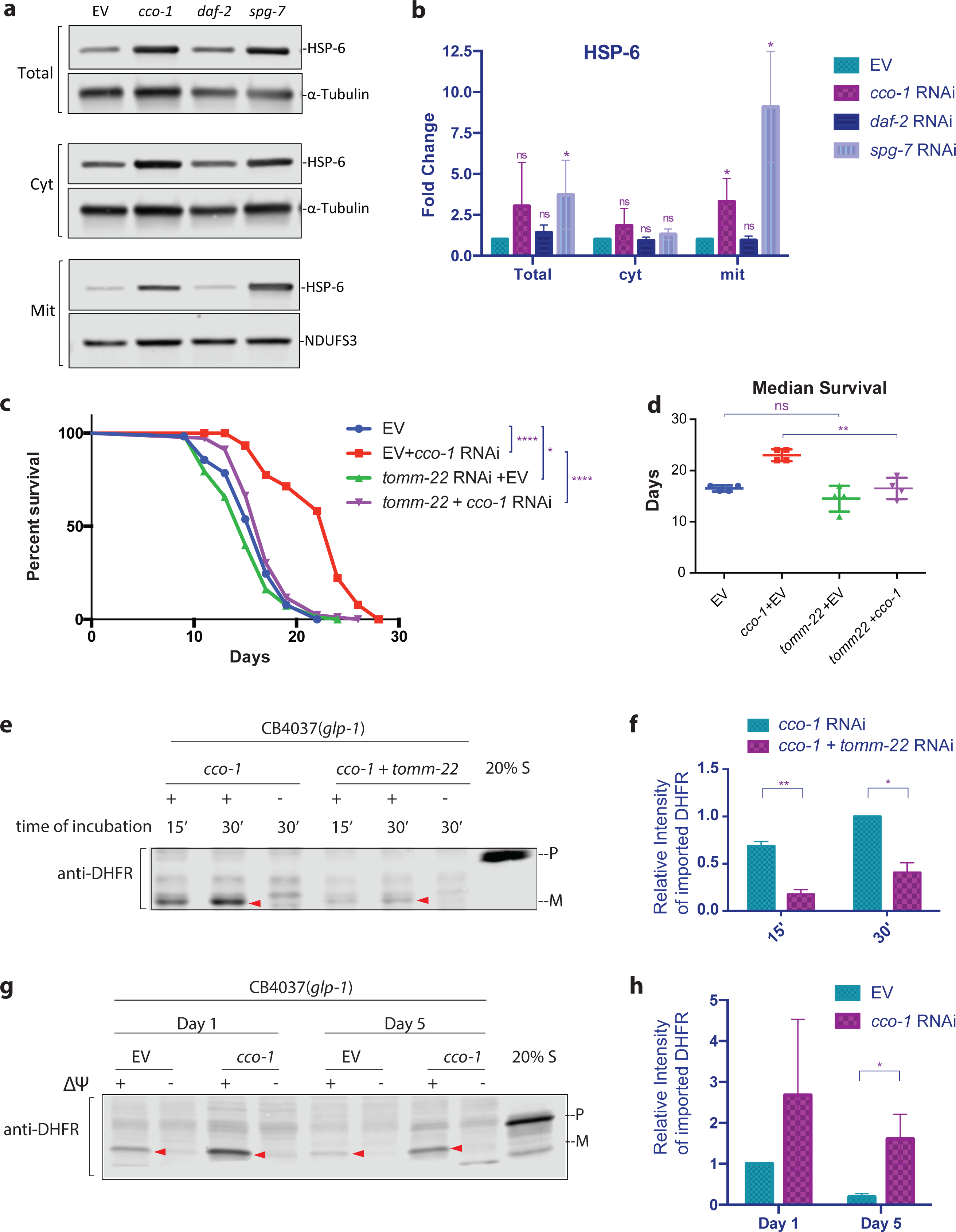
Import machinery is required for UPR^mt^-dependent lifespan extension, but UPR^mt^ does not prevent the age-associated decline of import. **a**, Subcellular fractionation and western blot of mitochondrial chaperone HSP-6. *glp-1* animals were grown at 25°C on bacteria expressing *cco-1, spg-7*, or *daf-2* dsRNA (each was diluted to 1:1 ratio with bacteria containing the empty RNAi vector alone) from hatching until the first day of adulthood. Control worms were grown on bacteria containing the empty RNAi vector alone. Different fractions were separated via differential centrifugation. **b**, Signal intensity was normalized to alpha-tubulin (total and cytosolic fraction) or NDUFS3 (mitochondrial fraction). HSP-6 level in each fraction was compared to that of the respective empty RNAi vector control and analyzed with unpaired student’s t-test. The graph is presented as mean ± SD of four biological repeats (*P<0.05, **P<0.01). **c**, Adult lifespans of CF512 *fer-15(b26ts); fem-1(hc17)* animals grown on bacteria expressing dsRNA from the time of hatching. Blue lines, the lifespan of animals grown on control bacteria containing the RNAi vector alone; red lines, the lifespan of animals grown on bacteria expressing the dsRNA of *cco-1*; purple lines, the lifespan of animals grown on bacteria expressing the dsRNA of *cco-1* and the dsRNA of *tomm-22*; green lines, the lifespan of animals grown on bacteria expressing the dsRNA of *tomm-22*. Log-rank (Mantel-Cox) method was used to determine the significant differences (*P<0.05, ****P<0.0001). **d**, Median lifespans with four replications are plotted and analyzed with unpaired student’s t-test. **P<0.01. **e** and **f**, *glp-1(e2141ts*) worms were synchronized and grown at 25°C on bacteria expressing dsRNA from the time of hatching. Mitochondria were isolated on day 1 of adulthood and subjected to import assay followed by western blot analysis. Import efficiency was quantified by measuring the mature imported protein as detected by DHFR antibody, followed by analysis with unpaired student’s t-test (**f**). **g** and **h**, *glp-1(e2141ts*) worms were synchronized in two batches and grown at 25°C. When the two batches reached day 1 and 5 of adulthood, respectively, mitochondria were isolated in parallel and subjected to import assay with 30 minutes incubation time followed by western blot analysis. Import efficiency was quantified by measuring the mature imported protein as detected by the DHFR antibody, followed by analysis with unpaired student’s t-test (**h**). All graphs are presented as mean ± SD of two to four biological repeats (*P<0.05, **P<0.01). Arrowheads: mature (imported) DHFR with the MTS cleaved off.

The next question we asked was whether the improvement of import is a necessary element for UPR^mt^-induced longevity. To test this, we used double RNAi to knockdown import activity in long-lived *cco-1*-deficient animals. We found that when worms were treated with *tomm-22* RNAi simultaneously with *cco-1* RNAi, both the enhancement of import (Fig. 4e, f, Extended Data Fig. 4b) and the extension of lifespan was largely suppressed, whereas *tomm-22* RNAi had a minimal effect on lifespan in wild-type animals (Fig. 4c, d). Similarly, treating worms with *timm-17* RNAi also partially suppressed the lifespan extension (Extended Data Fig. 4c). Taken together, these results indicate that intact import machinery is essential for UPR^mt^-induced lifespan extension.

### Induction of the UPR^mt^ maintains mitochondrial import during aging

Intrigued by the finding that enhanced import efficiency was required for UPR^mt^-mediated longevity, we asked how import efficiency might play a role in normal aging. As organisms age mitochondria gradually depolarize and the membrane potential, the major driving force of import, declines ^22^. We tested import efficiency across mitochondria isolated from aging cohorts of *C. elegans*. Indeed, we confirmed that mitochondrial import declines dramatically as the animals age (Extended Data Fig. 4d-f). Three age groups (Days 1, 5, and 9) were chosen to represent worms in the process of aging. We observed a more than one-half reduction in the import efficiency of su9-DHFR from Day 1 to Day 5, and no further decline from Day 5 to Day 9. This finding suggests a catastrophic loss in import capacity early in the aging process. Similar age-associated decline in import was observed in both *glp-1(e2141ts)* worms, which lack a germline (Fig. 4g, h, Extended Data Fig. g), and CF512 *fer-15(b26ts); fem-1(hc17)* worms, which have intact female germline (Extended Data Fig. 4d-f), suggesting that import in somatic tissue and germline are both affected by aging.

Treating worms with RNAi against *cco-1* delayed the age-associated decline of import. For example, import remained significant on day 5 in *cco-1* RNAi treated worms, whereas import in mock-RNAi control worms was barely detectable (Fig. 4g, h, Extended Data Fig. 4g). The transcriptional level of TIM/TOM import machinery also remained higher in older worms under *cco-1* RNAi treatment (Extended Data Fig. 4h).

As lifespan extension is a common effect of UPR^mt^ activation, it is intriguing to know whether the enhancement in mitochondrial import capacity we observed is a secondary effect of delayed aging or more specific to UPR^mt^ induced forms of longevity. *glp-1(e2141ts)* mutant worms, in which germline deficiency induces longevity, display highly compromised import capacity (Extended Data Fig. 1f-n), thus arguing against a causative effect of delayed aging on increasing mitochondrial import. We also tested another major pathway that regulates lifespan, the insulin/IGF1 signaling pathway mediated by the insulin/IGF1 receptor, *daf-2* in worms. We found that import capacity in day 1 adult worms was not affected by either *daf-2* RNAi or the *daf-2(e1370)* mutation (Extended Data Fig. 2j-l and m-o), despite the dramatic increase in lifespan in these strains. In addition, upon knockdown of *daf-2*, we did not observe an increase in import machinery, either at the transcription level (Extended Data Fig. 2i) or protein level (Fig. 2f, g). Similarly, the mitochondrial chaperone HSP-6 is not enhanced with *daf-2* RNAi (Fig. 4a, b). Taken together, these findings indicate that the enhancement of import in somatic cells is unique to UPR^mt^ activation, and the aging process itself reduces import that can be combated by UPR^mt^ activation.

## Discussion

Metazoans have evolved various defense mechanisms to protect themselves against the detrimental consequences of stress and aging. Many of the stress responsive mechanisms require altering the composition of their proteomes. This remodeling often includes enhancing the networks of stress responsive proteins and chaperones, which are targeted for specific subcellular compartments or organelles that are stressed.

It has been proposed that the alteration of mitochondrial import plays a role in the induction of mitochondrial unfolded protein response ^7^. However, the regulation of mitochondrial import upon stress has not been investigated in depth. In this work, we revealed the augmentation of mitochondrial protein import as a downstream effect of the mitochondrial stress response. This upregulation of import is specifically associated with the induction of UPR^mt^, instead of being a generic secondary effect of delayed aging or prolonged lifespan. We did not observe upregulation of the mitochondrial membrane potential that correlates with the import competency. In fact, we found that in spite of decreased membrane potential, the UPR^mt^ resulted in increased import activity. Taken together, our findings indicate that the enhancement of import is mediated by transcriptional regulation of the mitochondrial import machinery, and our *in vitro* biochemical assays reveal increased import competency of these mitochondria. Intriguingly, the efficiency of mitochondrial import serves as an active mechanism of increased longevity upon the activation of the UPR^mt^.

It is of great importance to establish the UPR^mt^ activation paradigm in mammalian cells and thereby analyze its impact on the mitochondrial import. Notably, loss of Tfam in T cells have reduced mitochondrial respiration and lower ETC components, but the levels of TOM20 seem to be elevated ^35^. As lowered levels of ETC components would presumably induce UPR^mt^, this observation in mammalian cells is consistent with our finding that import machinery is upregulated upon UPR^mt^ activation.

It was previously proposed that mitochondrial import deteriorates upon mitochondrial stress and thereby excludes ATFS-1 from mitochondria, allowing it to enter the nucleus to induce the expression of downstream stress response genes ^7^. Surprisingly, we found that the UPR^mt^-dependent upregulation of import is not only true for general import, but also the case for ATFS-1, the import-deficiency-dependent messenger of stress. Our findings raise the question of if and how the UPR^mt^ is maintained upon upregulation of the UPR^mt^. It was recently revealed that mitochondrial stress during larval development induces chromatin changes that are perpetuated into adulthood and make up a critical part of the UPR^mt 8,9^. Accordingly, UPR^mt^ induced by a transient deficiency in import may be sufficient to self-sustain the downstream effects, including the prolonged upregulation of import. In fact, it was found that the expression of ATFS-1 itself is upregulated upon induction of the UPR^mt 7^, suggesting that once the UPR^mt^ is activated, nuclear ATFS-1 might be kept at a higher level despite the recovery of mitochondrial import efficiency.

Notably, mitochondria exhibit a high level of heterogeneity within cells ^36^. It is conceivable that among the large population of mitochondria within a cell, some might remain at a low-import status and constantly send stress signal to the nucleus, whereas the rescuing proteins, once made, are sent to relatively healthy sub-population of mitochondria, or are used in the genesis and assembly of a new cohort of mitochondria. Therefore, the higher level of import efficiency upon UPR^mt^ in the *in vitro* import assay may be due to a fraction of mitochondria being healthier and more resilient due to UPR^mt^ activation and could be better preserved in the extraction process. In the future, it will be imperative to monitor the mitochondrial import of individual mitochondria *in vivo* during the aging process and under conditions of UPR^mt^ induction.

## Figure Legends

**Extended Data Figure 1.**
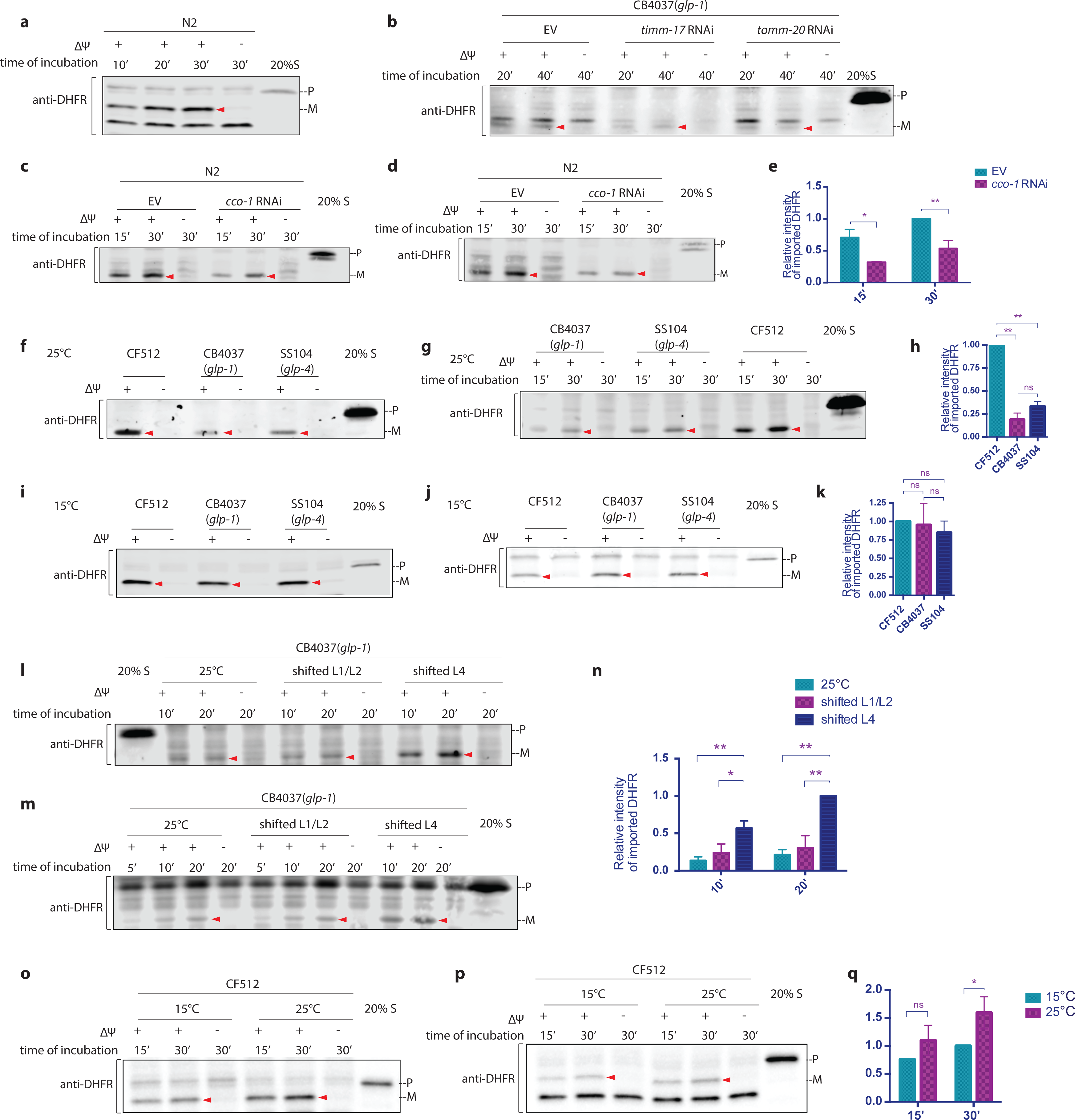
Germline-deficiency leads to the reduction of import competency; Somatic and germ cells differ in their import capacity. **a**, A biological replicate of Fig. **1b**. **b**, A biological replicate of Fig. **1d**. **c**-**e**, N2 wild type worms were synchronized and treated with RNAi against *cco-1.* Mitochondria were isolated at day 1 of adulthood and subjected to the import assay (**c** and **d** are biological replicates)**. f-k**, Temperature-sensitive germline-deficient *glp-1(e2141ts)* and *glp-4(bn2ts)* and spermatogenesis mutant strain CF512, were grown at the restrictive, 25°C (**f**), or permissive temperature, 15°C (**i**). Mitochondria were isolated at day 1 of adulthood and subjected to the import assay with 30 minutes incubation time (**f** and **g** are biological replicates, and **i** and **j** are biological replicates). **l-n**, *glp-1(e2141ts)* mutant worms were synchronized and shifted at different developmental stages to the restrictive temperature. Mitochondria were then isolated and subjected to the import assay (**l** and **m** are biological replicates). **o-q**, CF512 worms were synchronized and raised at 15°C or 25°C. Mitochondria were then isolated and subjected to the import assay (**o** and **p** are biological replicates). The efficiency of mitochondrial import was quantified by measuring the mature imported protein as detected by the DHFR antibody, followed by analysis with unpaired student’s t-test (**e, h, k, n, q)**. All graphs are presented as mean ± SD of two to four biological repeats. *P<0.05, **P<0.01. Arrowheads: mature (imported) DHFR with the MTS cleaved off.

**Extended Data Figure 2.**
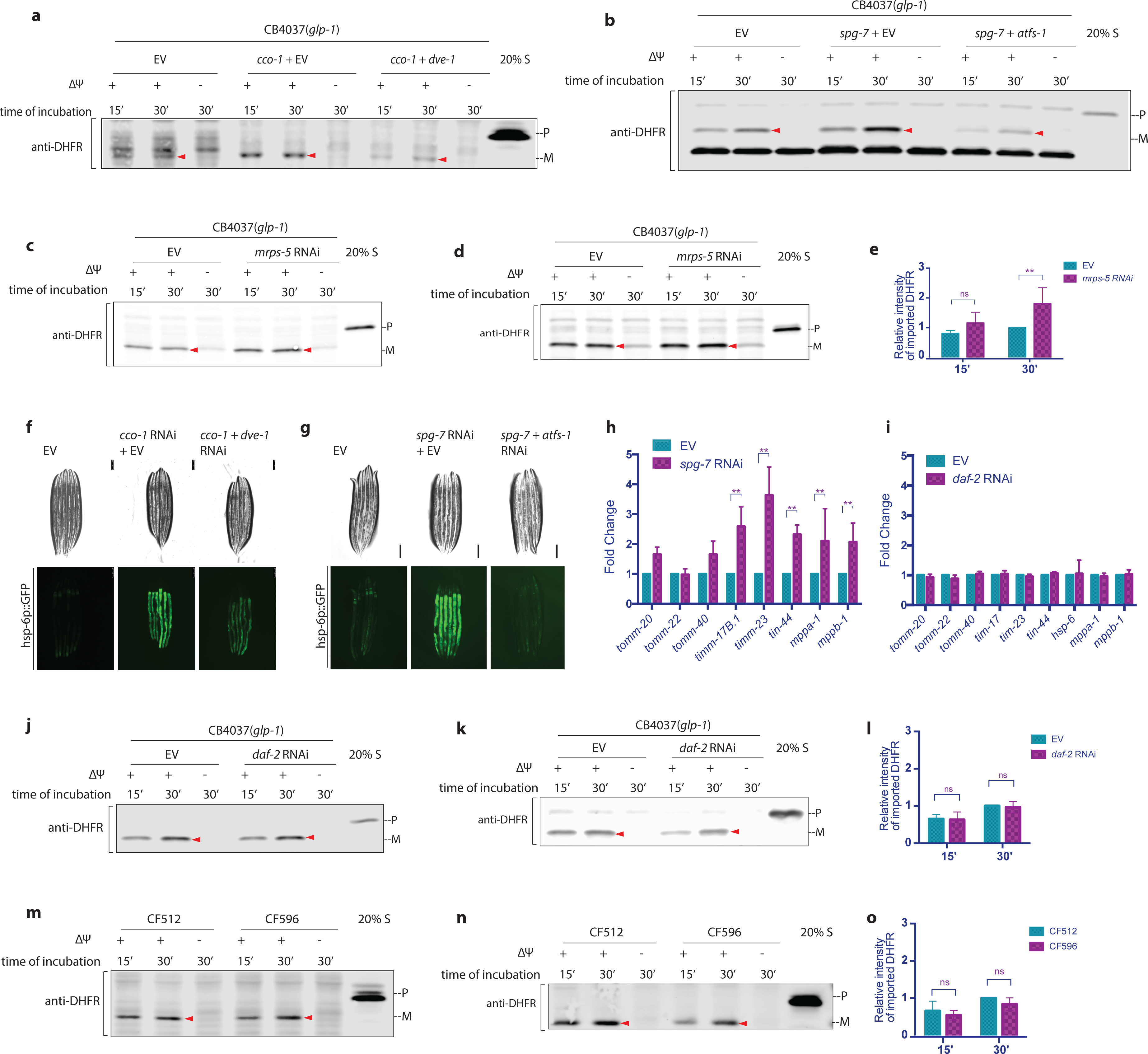
The UPR^mt^ promotes mitochondrial import. **a**, A biological replicate of Fig. **2a. b**, A biological replicate of Fig. **2c**. **c-e**, *glp-1(e2141ts*) mutant animals were grown at 25°C on bacteria expressing *mrps-5* dsRNA (20% diluted with bacteria containing the empty RNAi vector alone) from hatching until the first day of adulthood. Mitochondria were isolated on day 1 of adulthood and subjected to import assay followed by western blot analysis (**c** and **d** are biological replicates). **f** and **g**, To induce UPR^mt^ during development, *hsp-6p*::GFP animals were grown at 25°C on bacteria expressing *cco-1* dsRNA (**f**) or *spg-7* dsRNA (**g**) (diluted to 1:1 ratio with bacteria containing the empty RNAi vector alone) from the time of hatching until the first day of adulthood. To suppress UPR^mt^, animals were treated with double RNAi (1:1 mixture of bacteria replacing the empty RNAi vector with *dve-1* dsRNA (**f**) or *atfs-1* dsRNA (**g**). Control worms were grown on bacteria containing empty vector alone. **h** and **i**, *glp-1(e2141ts*) animals were grown at 25°C on bacteria expressing *spg-7* dsRNA (**h**) or *daf-2* dsRNA (**i**) (diluted to 1:1 ratio with bacteria containing the empty RNAi vector alone) from hatching until the first of adulthood. RNA was isolated on day 1 of adulthood, and qPCR analysis was performed. Expression was normalized against three housekeeping genes. **j**-**l**, *glp-1(e2141ts*) animals were grown at 25°C on bacteria expressing *daf-2* dsRNA (diluted to 1:1 ratio with bacteria containing the empty RNAi vector alone) from hatching until the first day of adulthood. Mitochondria were isolated on day 1 of adulthood and subjected to import assay followed by western blot analysis (**j** and **k** are biological replicates). **m-o**, CF512 *fer-15(b26) II; fem-1(hc17ts) I* and CF596 *daf-2(mu150) III; fer-15(b26); fem-1(hc17ts)* worms were grown at 20°C until larval stage L2 and then transferred to 25°C. Mitochondria were isolated on day 1 of adulthood and subjected to import assay followed by western blot analysis (**m** and **n** are biological replicates). Import efficiency was quantified by measuring the mature imported protein as detected by the DHFR antibody, followed by analysis with unpaired student’s t-test (**e, l**, and **o)**. All graphs are presented as mean ± SD of three or more biological repeats. *P<0.05, **P<0.01. Arrowheads: mature (imported) DHFR with the MTS cleaved off.

**Extended Data Figure 3.**
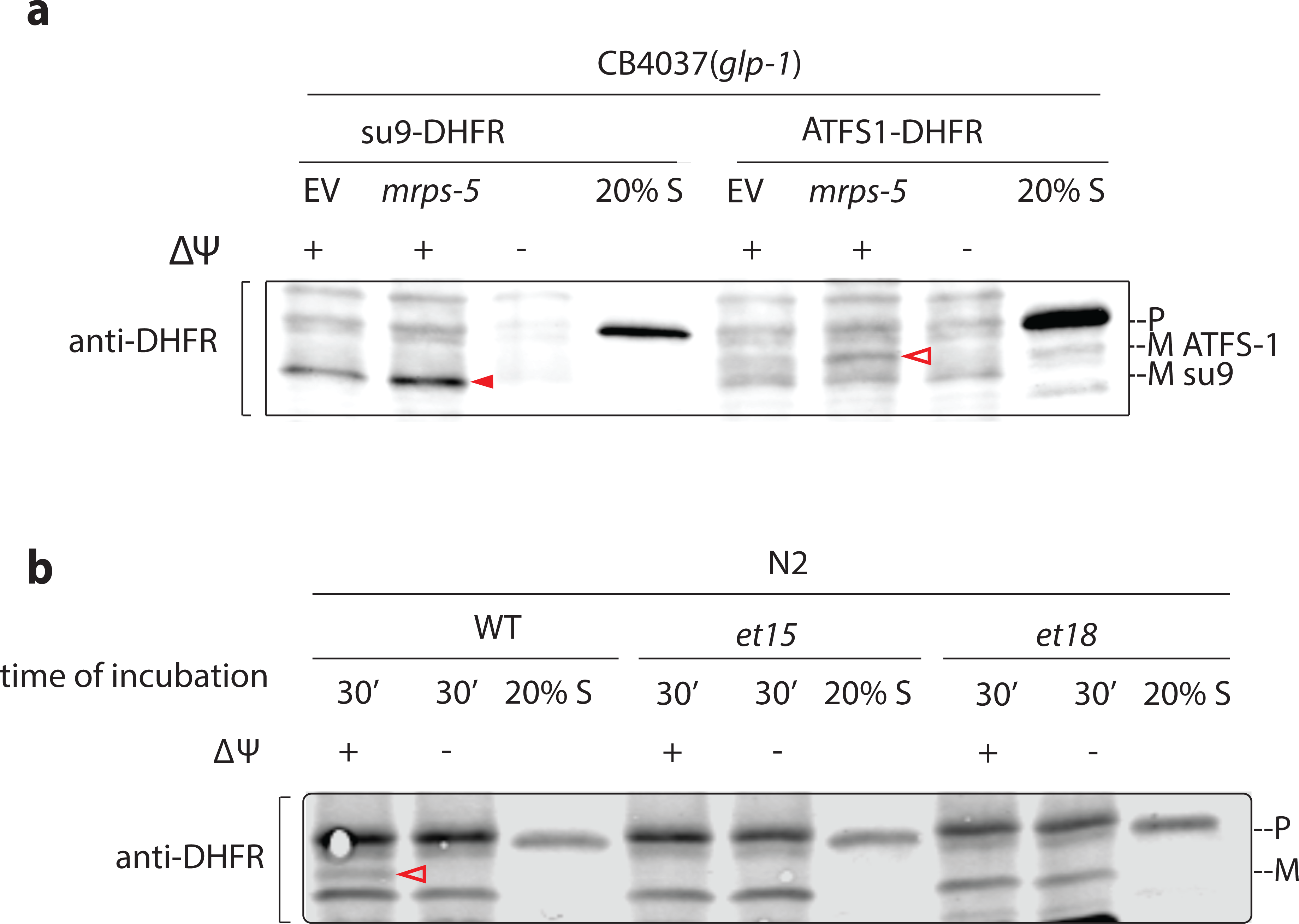
**a**, A biological replicate of Fig. **3a**. **b**, A biological replicate of Fig. 3d. Incubation time: 30 minutes. Solid arrowheads: mature (imported) su9-DHFR with the su9 MTS cleaved off. Open arrowheads: mature (imported) ATFS1-DHFR with the MTS of ATFS-1 cleaved off.

**Extended Data Figure 4.**
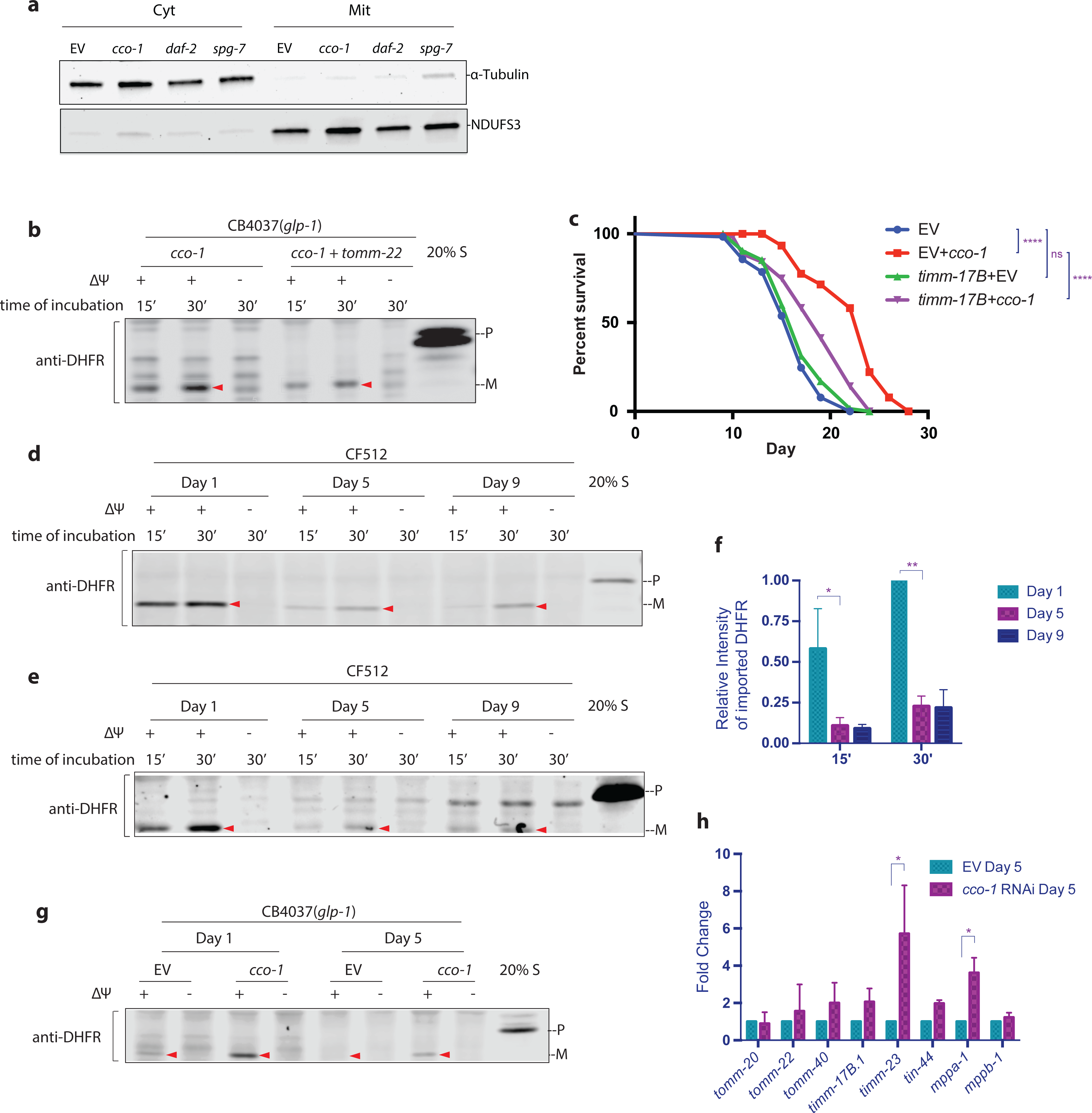
Import machinery is required for UPR^mt^-dependent lifespan extension. **a**, The enrichment of subcellular extracts was confirmed with western blot analysis using the subcellular fraction-specific antibodies against NDUFS3 (mitochondria) and alpha-tubulin (cytosol). **b**, A biological replicate of Fig. **4e**. **c**, Adult lifespan of CF512 animals grown on bacteria expressing dsRNA from the time of hatching. Blue line, the lifespan of animals grown on control bacteria containing the RNAi vector alone; red line, the lifespan of animals grown on bacteria expressing the dsRNA of *cco-1*; purple line, the lifespan of animals grown on bacteria expressing the dsRNA of *cco-1* and the dsRNA of *tim-17*; green line, the lifespan of animals grown on bacteria expressing the dsRNA of *tim-17*. Log-rank (Mantel-Cox) method was used to determine the significant differences (****P<0.0001). **d-f**, CF512 worms were synchronized in three batches and grown at 25°C. When the three batches reached day 1, 5, and 9 of adulthood, respectively, mitochondria were isolated in parallel and subjected to import assay followed by western blot analysis (**c** and **d** are biological replicates). Import efficiency was quantified by measuring the mature imported protein as detected by DHFR antibody, followed by analysis with unpaired student’s t-test (**e**). **g**, A biological replicate of Fig. **4g**. **h**, *glp-1(e2141ts*) worms were synchronized and grown at 25°C on bacteria expressing dsRNA from the time of hatching. RNA was isolated on day 5 of adulthood, and qPCR analysis was performed. Expression was normalized against three housekeeping genes. All graphs are presented as mean ± SD of two to three biological repeats (*P<0.05, **P<0.01). Arrowheads: mature (imported) DHFR with the MTS cleaved off.

## Acknowledgment

We thank the Caenorhabditis Genetic Center and Shohei Mitani of the National BioResource Project for providing strains.

This research was supported by NIH F32 (5F32AG051353-03), NIH R01 (R37AG024365 and R01ES021667), University of Texas Health Science Center at Houston (37516-12002) and the Rising STARs program of UT systems (26532). This work used the Vincent J. Coates Genomics Sequencing Laboratory at UC Berkeley, supported by NIH S10 Instrumentation Grants S10RR029668 and S10RR027303.

We thank Dr. Stephan Rolland from Ludwig Maximilian University of Munich for sharing a sequence-verified *timm-23* RNAi strain.

We thank Dr. Ryo Higuchi-Sanabria, Dr. Hope Henderson, and the Dillin Lab for comments that greatly improved the manuscript.

We would also like to show our gratitude to Dr. Ye Tian (Dillin Lab alumni, Institute of Genetics and Developmental Biology, CAS), for sharing her insight with us during the course of this research.

## Methods

### Strains

CB4037 *glp-1(e2141)* III, SS104 *glp-4(bn2) I*, CF512 *fer-15(b26) II, fem-1(hc17ts)* IV, CF596 *daf-2(mu150) III; fer-15(b26); fem-1(hc17ts)*, SJ4100 *(zcIs13[hsp-6p::gfp])*, N2 wild-type strains were obtained from the Caenorhabditis Genetics Center (Minneapolis, MN).

### RNAi Feeding

Worms were grown from the hatch on HT115 *Escherichia coli* containing an empty vector control or expressing double-stranded RNA. RNAi strains were from the Vidal library if present or the Ahringer library if absent from the Vidal library.

### Import assay

pGEM4-su9(1-69)-DHFR plasmid was a gift from Dr. Thomas Langer ^37^. Fusion protein su9-DHFR was transcribed, translated, and biotinylated in a single reaction with the TnT® SP6 Quick Coupled Transcription/Translation System (Promega, L2080) and Transcend™ tRNA (Promega L5016). Mitochondria extraction was made from synchronized worms at the designated age in mitochondria extraction buffer (5 mM Tris-HCl pH 7.4, 210 mM mannitol, 70 mM sucrose, 0.1 mM EDTA). Protease Inhibitor (Protease Inhibitor Cocktail Set III, EDTA-Free, Calbiochem 539134) was used at 1:1000). Worms were mechanically homogenized with Dura-Grind™ Stainless Steel Dounce Tissue Grinder (Wheaton 357572), and mitochondria were isolated via differential centrifugation. Mitochondria pellets were resuspended in buffer C (20mM potassium HEPES, 0.6M sorbitol). Protein concentration was measured using BCA Protein Assay Kit (Pierce 23225), and the same amount is used in each import reaction.

Mitochondrial import assay was performed as previously described ^14^ with some modification. The biotinylated protein was incubated with 50ug fresh mitochondria extraction in import buffer containing ATP regeneration system (Creatine kinase (Roche 10127566001), Creatine Phosphate (Sigma-Aldrich 10621714001) for 10 to 45 minutes at 25°C with gentle shaking. Import assay with a single time point was performed with a 30-minutes incubation time unless otherwise noted. Membrane potential (ΔΨ) was disrupted with valinomycin in control. Mitochondria were subsequently treated with proteinase K to degrade preproteins that are attached to the surface of the mitochondria. Mitochondria were spun down and resuspended in mitochondria extraction buffer. SDS (6×) loading buffer was added to each sample. Samples were heated at 95°C for 5 min and resolved by NuPAGE Bis-Tris mini gels, followed by western blot with DHFR antibody and streptavidin. Import efficiency was quantified by measuring the mature imported protein as detected by the DHFR antibody in ImageStudio (LiCor). Signal intensity was normalized against the signal intensity of control treatment with 30-minute incubation time unless otherwise noted. Data were analyzed using unpaired t-test with Prism (GraphPad4).

### Subcellular Fractionation

Synchronized worms were lysed, and mitochondria were isolated as previously described ^38^. Supernatant before and after the centrifugation for mitochondria are kept as total and cytosolic portion, respectively.

### qPCR

Total RNA was harvested from worms at the early adult Day 1 stage using TRIzol® LS Reagent (Life Technologies). After freezing and thawing three times, RNA was purified on RNeasy mini columns (QIAGEN), and cDNA was synthesized using the QuantiTect Reverse Transcription kit (QIAGEN). SybrGreen quantitative RT-PCR experiments were performed as described in the manual using QuantStudio™ 6 Flex Real-Time PCR System. Internal controls utilized a geometric mean of *cdc-42, pmp-3*, and *Y45F10D.4*. Experiments were repeated three times. Primers used for qPCR are listed below.

hsp-6 forward 5’-CAAACTCCTGTGTCAGTATCATGGAAGG-3’

hsp-6 reverse 5’-GCTGGCTTTGACAATCTTGTATGGAACG-3’

tomm-20 forward 5’-CGGCTACTGCATTTACTTCGA-3’

tomm-20 reverse 5’-TCATTGCCTGCTGCAGCTGGA-3’

tomm-22 forward 5’-CGACTTCGTTCAGCAGTTCAT-3’

tomm-22 reverse 5’-GCGATCAATGACGTTGTAGATA-3’

tomm-40 forward 5’-AGCTCGTGATGTCTTCCCAAC-3’

tomm-40 reverse 5’-TCCAAATCGGTATCCGGTGTT-3’

timm-17B.1 forward 5’-GATTGTTGTCTTGTCGCCATCC-3’

timm-17B.1 reverse 5’-ATCACCTTTGGTCCTGAACGG-3’

timm-23 forward 5’-AGTGCCGGAATGAACTTCTC-3’

timm-23 reverse 5’-GTTGATCCAAGGCGAGGAC-3’

tin-44 forward 5’-GGGATACGATTAACTCGGACA-3’

tin-44 reverse 5’-CTGCATTCGAGCTTTCAACTG-3’

mppa-1 forward 5’-CGATTTTGTGACTGTTGGCGT-3’

mppa-1 reverse 5’-GCTTGAGAACGATTCCGATGA-3’

mppb-1 forward 5’-GCACAAGTTCAGCCGAAATCA-3’

mppb-1 reverse 5’-TTCTCATTCTCGTAGCGACTG-3’

cdc-42 forward 5’-AGGAACGTCTTCCTTGTCTCC -3’

cdc-42 reverse 5’- GGACATAGAAAGAAAAACACAGTCAC -3’

pmp-3 forward 5’- CGGTGTTAAAACTCACTGGAGA -3’

pmp-3 reverse 5’- TCGTGAAGTTCCATAACACGA -3’

Y45F10D.4 forward 5’- AAGCGTCGGAACAGGAATC -3’

Y45F10D.4 reverse 5’- TTTTTCCGTTATCGTCGACTC -3’

### Antibodies

A polyclonal rabbit antibody to TIMM-23 was generated against the synthesized polypeptide of amino acid 95-230, and affinity-purified (ABClonal Science, Inc.). Other antibodies and probes used for western blot were as follows: anti-DHFR antibody (Sigma-Aldrich D1067), anti-HSP-6 antibody (Enzo Life Science ADI-SPS-825); and anti-NDUF3 [17D95] antibody (Abcam ab14711); IRDye® 680CW Donkey anti-Mouse IgG (H + L) (LI-COR 926-68072); IRDye 680LT Donkey anti-Rabbit IgG (H + L), (LI-COR 926-68023).

### TMRE Staining

TMRE staining was performed according to the previous study ^39^. TMRE was dissolved in DMSO at a concentration of 50µM and added into fresh bacteria culture at a final concentration of 0.1µM before seeding the plates. Worms were synchronized by egg bleach and grown on *E. coli* HT115 for RNAi from the hatch and transferred to RNAi plates containing TMRE at the L3/L4 stage. Worms were imaged after growing overnight on TMRE plates. TMRE staining was quantified with ImageJ.

CCCP is dissolved in DMSO at a concentration of 10mM and added into bacteria culture at a final concentration of 50µM before seeding the plates.

### Lifespan Analysis

Lifespan experiments were performed with CF512 worms at 25°C as previously described ^18^. Worms were synchronized by egg bleach and grown on *E. coli* HT115 for RNAi from the hatch. Worms were scored every second day. Prism 6 software was used for statistical analysis. Log-rank (Mantel-Cox) method was used to determine the significant difference.

